# Cholinergic regulation of prolonged alcohol withdrawal differs by sex

**DOI:** 10.64898/2025.12.12.693968

**Authors:** NR Payne, SM Scott, M Scalf, RF Dobbelmann, S Engel, AM Lee

## Abstract

Acute alcohol withdrawal encompasses somatic withdrawal signs and increased negative affect. In prolonged alcohol withdrawal the somatic withdrawal signs have resolved but the increased negative affect persists. We investigated acute and prolonged alcohol withdrawal after 9 daily injections of 2.5 g/kg alcohol plus 4-methypyrazole, an alcohol dehydrogenase inhibitor, in male and female C57BL/6J mice, and examined whether nicotinic acetylcholine receptor (nAChR) drugs could attenuate withdrawal-induced negative affect. Male mice showed changing somatic withdrawal signs over time and negative affect that persisted at least 21 days into withdrawal. Pre-treatment with mecamylamine, a non-specific nAChR antagonist, or varenicline, a nAChR partial agonist, reduced withdrawal-induced anxiety- and compulsive-like behavior in the marble-burying test during prolonged withdrawal. In contrast, female mice did not exhibit somatic withdrawal signs or anxiety- or compulsive-like behaviors. Instead, female mice showed a deficit in social interaction that was not attenuated by mecamylamine. Alcohol clearance and sedation were not different between sexes, indicating that differences in withdrawal signs and negative affect are not confounded by differences in alcohol metabolism. These findings suggest that cholinergic drugs may be a promising therapeutic for withdrawal-induced negative affect in male, but not female, mice.

## Introduction

There are numerous sex differences in the effects of alcohol and the course of alcohol use disorder (AUD) in humans. Women consume as much alcohol as men but experience alcohol-related harms at lower doses and over shorter time periods compared with men [1]. Women are also more likely to experience hangovers and blackouts compared with men when at the same blood alcohol concentration [2]. Lastly, women are also less likely to be treated for AUD compared with men [3].

Alcohol withdrawal in humans can be categorized into acute and prolonged withdrawal. Acute withdrawal encompasses somatic withdrawal signs and negative affective changes that occur in the first days to weeks of abstinence, whereas prolonged withdrawal describes a time frame of months or years into abstinence when somatic withdrawal signs have long resolved but negative affect persists. Prolonged withdrawal symptoms of AUD had been described in the 1950s and consisted of irritability, restlessness, depression, insomnia and distractibility, that “occur in waves of greater or lesser intensity” with most severe symptoms occurring in the first 6 months after abstinence [4]. This description highlights a constellation of negative affective symptoms with a temporal component. Subsequent studies and meta-analyses have included additional symptoms such as negative mood and anxiety, anhedonia, cognitive impairment and sleep disturbances in prolonged withdrawal [5–7]. Importantly, these persistent behavioral disturbances in prolonged withdrawal are a critical risk factor for relapse back to alcohol use [8].

Most of the studies describing prolonged negative affect in alcohol withdrawal have primarily collected data from men and how prolonged alcohol withdrawal presents in women has not been well described. Meta-analyses of past work have shown sex differences in the clinical presentation of alcohol withdrawal as men tend to exhibit more withdrawal signs and tremors compared with women; however, women are vastly underrepresented in the data and the majority of the data was collected during the acute withdrawal phase and not during prolonged withdrawal [9, 10].

In pre-clinical research, mice also exhibit acute and prolonged withdrawal from alcohol, and show increased negative affect during prolonged alcohol withdrawal. Mice show increased anxiety- and compulsive-like behavior, increased anhedonia and reduced exploration activity [11]. Similar to human studies, the majority of the data has been collected from male animals, and sex differences in alcohol withdrawal have been reported as female mice tend to exhibit reduced alcohol withdrawal signs compared with males (for a review see [11]).

Current FDA-approved pharmacotherapies for AUD are disulfiram, acamprosate and naltrexone, and of these drugs naltrexone is the most well studied and is effective in reducing the number of drinking days, reducing craving and relapse risk [12]. However, additional drugs to treat AUD are needed to expand the available treatment options for patients, particularly in the prolonged withdrawal phase. One drug that has been in clinical trials for AUD is varenicline, a nicotinic acetylcholine receptor (nAChR) partial agonist that is FDA-approved for smoking cessation [13]. Varenicline has shown promise in decreasing alcohol consumption and reducing craving in clinical trials [14–18]. Varenicline has not been tested during the prolonged withdrawal phase.

The objective of this study was to examine acute and prolonged alcohol withdrawal in male and female C57BL/6J mice after chronic alcohol administration. We found that male mice exhibited different somatic withdrawal signs over time, and withdrawal-induced anxiety- and compulsive-like behavior that persisted at least 21 days into withdrawal. Mecamylamine, a non-specific nAChR antagonist, and varenicline both reduced withdrawal-induced behaviors in male mice. In contrast, female mice did not exhibit somatic withdrawal signs or anxiety- or compulsive-like withdrawal behaviors; however, female mice showed a deficit in social interaction that was not observed in male mice. Alcohol clearance was not different between male and female mice, indicating that differences in withdrawal signs and negative affect are not confounded by sex differences in metabolism.

## Methods

### Animals

Eight-to-twelve week old male and female C57BL/6J mice (The Jackson Laboratory) were group housed for a minimum of 5 days in our facility prior to starting experiments. Somatic withdrawal signs, elevated zero maze and the marble burying test were conducted in the same mice, and multiple cohorts of mice were run at different times. All behavioral tests were scored by personnel who were masked to the experimental group of the mice. All animal procedures and experiments were approved by the Institutional Animal Care and Use Committee (IACUC) at the University of Minnesota and were in accordance with recommendations of the ARRIVE guidelines (https://arriveguidelines.org).

### Alcohol and drug administration

Mice in the alcohol-treated groups received 9 daily *i.p.* injections of 2.5 g/kg alcohol (as 20% v/v alcohol in saline) plus 4-methylpyrazole (4MP, 9 mg/kg, cat #M1387, MilliporeSigma), an alcohol dehydrogenase inhibitor. Control-treated groups received 9 daily *i.p.* injections of saline + 9 mg/kg 4MP. In the behavioral tests examining the involvement of cholinergic activity, mecamylamine (3 mg/kg, Tocris Bioscience, #2843), varenicline (3 mg/kg, Tocris Bioscience, #3754) or saline were injected *i.p.* 15 minutes prior to behavioral testing.

### Somatic withdrawal signs

Mice were individually placed in a clean empty cage with bedding and were video recorded for 20 minutes. The videos were scored for withdrawal signs (grooming, digging, chewing, tremors/shakes and escape attempts). Behavioral data were then normalized to the control-treated group and expressed as a percentage of control. In the female mice, zero escape attempts were recorded at 24h and zero shakes were recorded at 7d. The time points were not recorded in the same group of mice and thus were analyzed without repeated measures for time.

### Marble Burying Test (MBT)

Mice were individually placed in clean rat-sized cages with bedding, with 20 marbles evenly placed in a grid, for 25 minutes. The number of marbles that were covered by 2/3^rd^ or more with bedding was noted. Mice used in experiments investigating the effect of mecamylamine or varenicline were only assessed in the MBT. Injections of mecamylamine, varenicline, or saline *i.p.* occurred 15 min prior to the MBT. In the experiments examining the effect of varenicline, varenicline or saline was administered to the same mice at both time points. Experiments using the same cohort of mice over multiple time points were analyzed using a repeated measure for time, whereas data that was collected in different cohorts for different time points were analyzed without repeated measures.

### Elevated zero maze (EZM)

Mice were placed on an elevated zero maze (Med Associates) for 5 minutes. The time spent in the open and closed arms were noted. The maze was cleaned in between each mouse with 70% alcohol to eliminate odors. The male data for the 7h and 24h time points, and the female data for the 24h and 7d time points were obtained with different cohorts of mice.

### Social Interaction Test (SIT)

Mice that were assessed for social interaction were tested in the SIT only. The SIT was conducted as previously described [19] under dim red lighting. The arena consisted of a translucent container with a wire mesh cup on one side. Each mouse underwent two, 3-minute trials. The first trial was in an empty arena, and the second trial had a novel mouse of the same sex under the wire mesh cup. The 3-minute trials were separated by less than 1 minute of time, and the trials were video recorded. The amount of time the mice spent investigating the wire mesh cup or in the opposite corners of the arena were noted. An interaction ratio was calculated that consists of the time interacting with novel mouse divided by the time spent in the corners for the novel mouse and empty arena trials. The female and male 24h and 7d SIT was performed in the same group of mice and thus analyzed with a repeated measure for time. The female data investigating the effect of mecamylamine on the SIT was obtained in a separate cohort of mice.

### Loss of Righting Reflex (LORR)

The LORR procedure was conducted as previously described [20]. All mice were injected with 4 g/kg alcohol (as 20% v/v in saline) *i.p.* and placed on their backs in a clean, empty cage (1^st^ LORR). The time to loss of the righting reflex was noted, and the righting reflex was considered regained when the mouse was able to right itself 3 times within 30 seconds. The total time without the righting reflex was calculated. The day after the 1^st^ LORR test, mice were assigned to an alcohol- or control-treated group and received 9 daily *i.p.* injections of 2.5 g/kg alcohol + 9 mg/kg 4MP or saline + 9 mg/kg 4MP, respectively. Mice underwent the same LORR test again 24h after the last injection (final LORR).

### Alcohol clearance

Male and female mice received either one *i.p.* injection of 2.5 g/kg alcohol + 9 mg/kg 4MP, or 9 daily injections of 2.5 g/kg alcohol + 9 mg/kg 4MP. Blood samples (∼15-20 μL) were collected from the facial vein at 15, 45, and 75 minutes following the last injection and were immediately centrifuged to separate plasma. One male mouse was sampled at 15 and 45 minutes only, and one female mouse was sampled at 45 and 75 minutes only. Plasma alcohol concentrations were measured using a colorimetric enzymatic assay (Neta Scientific, # CAYM-702260-96).

### Statistical Analyses

Data was normalized to the mean of the control-treated group and expressed as percent of control. Outliers were identified using the ROUT test. All data were analyzed with Prism 10 (GraphPad) and expressed as mean ± SEM. As prior research has shown numerous sex differences in alcohol withdrawal signs [11], we analyzed the sexes separately for withdrawal signs and negative affect. Our approach follows recent recommendations to analyze the sexes separately when the underlying relationship between variables is different between the sexes [21,22].

## Results

### Male somatic alcohol withdrawal signs change over time

Male C57BL/6J mice were given 9 daily injections of 2.5 g/kg alcohol plus 9 mg/kg 4-methylpyrazole (4MP), an alcohol dehydrogenase inhibitor that decreases the metabolism of alcohol. Control-treated mice were administered saline with 4MP. Somatic withdrawal signs were assessed at 7h, 24h and 7d after the last injection. We found that somatic alcohol withdrawal signs change over time in alcohol withdrawal in male mice. At 7h of withdrawal there was a significant interaction between the treatment group and the withdrawal sign (2-way ANOVA, Finteraction (4, 83)=15.71, *P*<0.0001; Fsign (4, 83)=15.71, *P*<0.0001; Ftx group (1, 83)=18.33, *P*<0.0001). A Sidak’s multiple comparisons test showed a significant increase in shaking episodes in the alcohol-treated mice compared with control-treated mice, with no differences in the other withdrawal signs (\*\*\*\**P*<0.0001 alcohol+4MP *vs.* control+4MP, **Fig 1A**). At 24h withdrawal, we found that alcohol-treated mice showed increased episodes of chewing behavior compared with the control-treated mice (2-way ANOVA, Finteraction (4, 68)=5.778, *P*=0.0005; Fsign (4, 68)=8.216, *P*<0.0001; Ftx group (1, 68)=9.397, *P*=0.003.

**Fig. 1.**
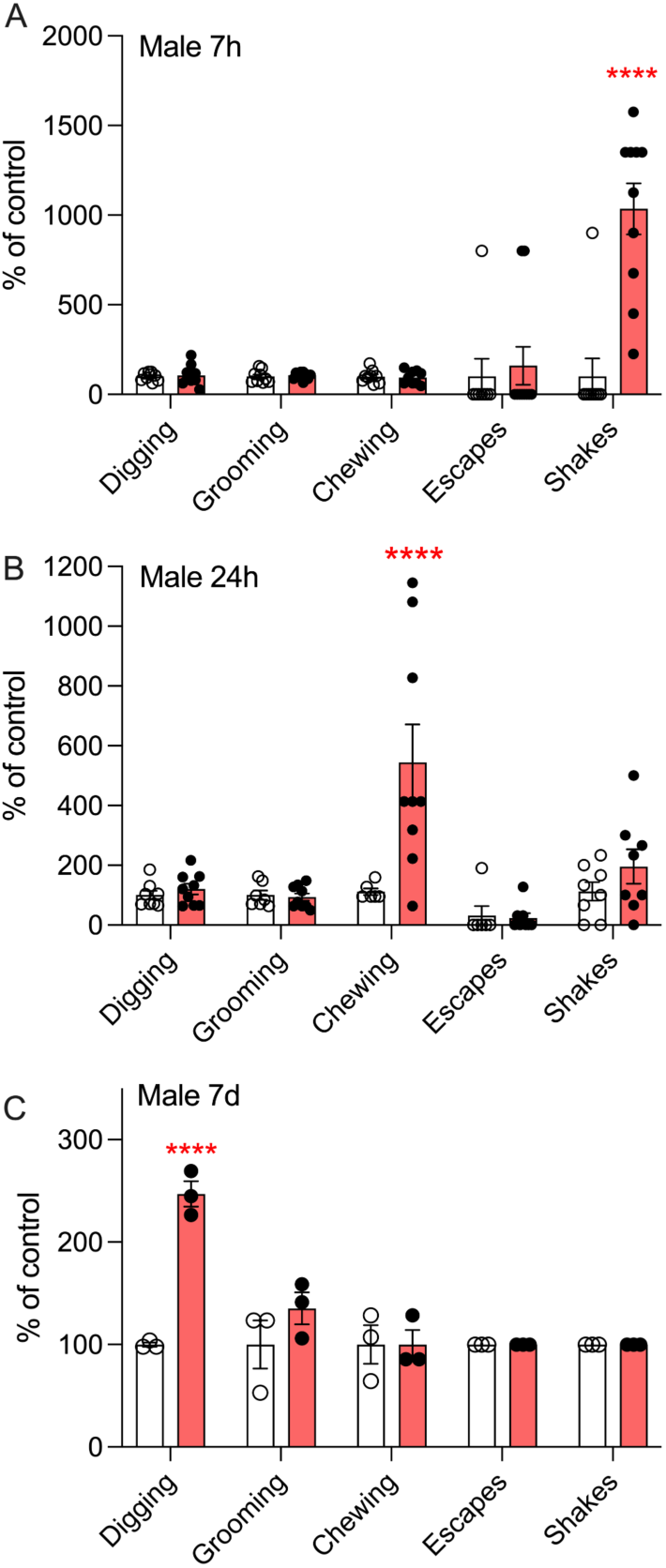
Alcohol withdrawal signs change over time in male C57BL/6J mice. Mice exhibit A) increased episodes of shakes at 7h withdrawal, B) increased episodes of chewing behavior at 24h withdrawal and C) increased episodes of digging behavior at 7d of withdrawal after the last alcohol exposure. Filled bars = alcohol withdrawn mice, clear bars = control mice. Sidak’s multiple comparisons test \*\*\*\**P*<0.0001 compared with control group. A, *n*=9-10 mice per group; B, *n*=8-9 per group; C, *n*=3 per group; data shown as mean ± SEM.

Sidak’s multiple comparisons test, \*\*\*\**P*<0.0001 alcohol+4MP *vs.* control+4MP, **Fig. 1B**). At 7d withdrawal, the alcohol-treated mice showed increased episodes of digging behavior compared with the control-treated mice (2-way ANOVA, Finteraction (4, 20)=13.36, *P*<0.0001; Fsign (4, 20)=13.36, *P*<0.0001; Ftx group (1, 20)=21.92, *P*=0.0001; Sidak’s multiple comparisons test, \*\*\*\**P*<0.0001 alcohol+4MP *vs.* control+4MP, **Fig. 1C**).

### Increased alcohol-withdrawal induced behaviors are attenuated by cholinergic drugs in male mice

Anxiety-like behavior was measured using the EZM at 24h and 7d after the last injection. There was a significant interaction between treatment group and time point (2-way ANOVA, Finteraction (1, 33)=5.196, *P*=0.03; Ftime (1, 33)=5.377, *P*=0.03; Fgroup (1, 33)=4.242, *P*=0.047), and a Sidak’s multiple comparison test showed a significant decrease in the time spent in the open arms in the alcohol-treated group at 24h withdrawal (\*\**P*=0.009, alcohol+4MP *vs.* control+4MP, **Fig. 2A**), indicating increased anxiety-like behavior in the alcohol-treated mice. There was no longer a difference in the percent time spent in the open arms at 7d withdrawal between groups.

**Fig. 2.**
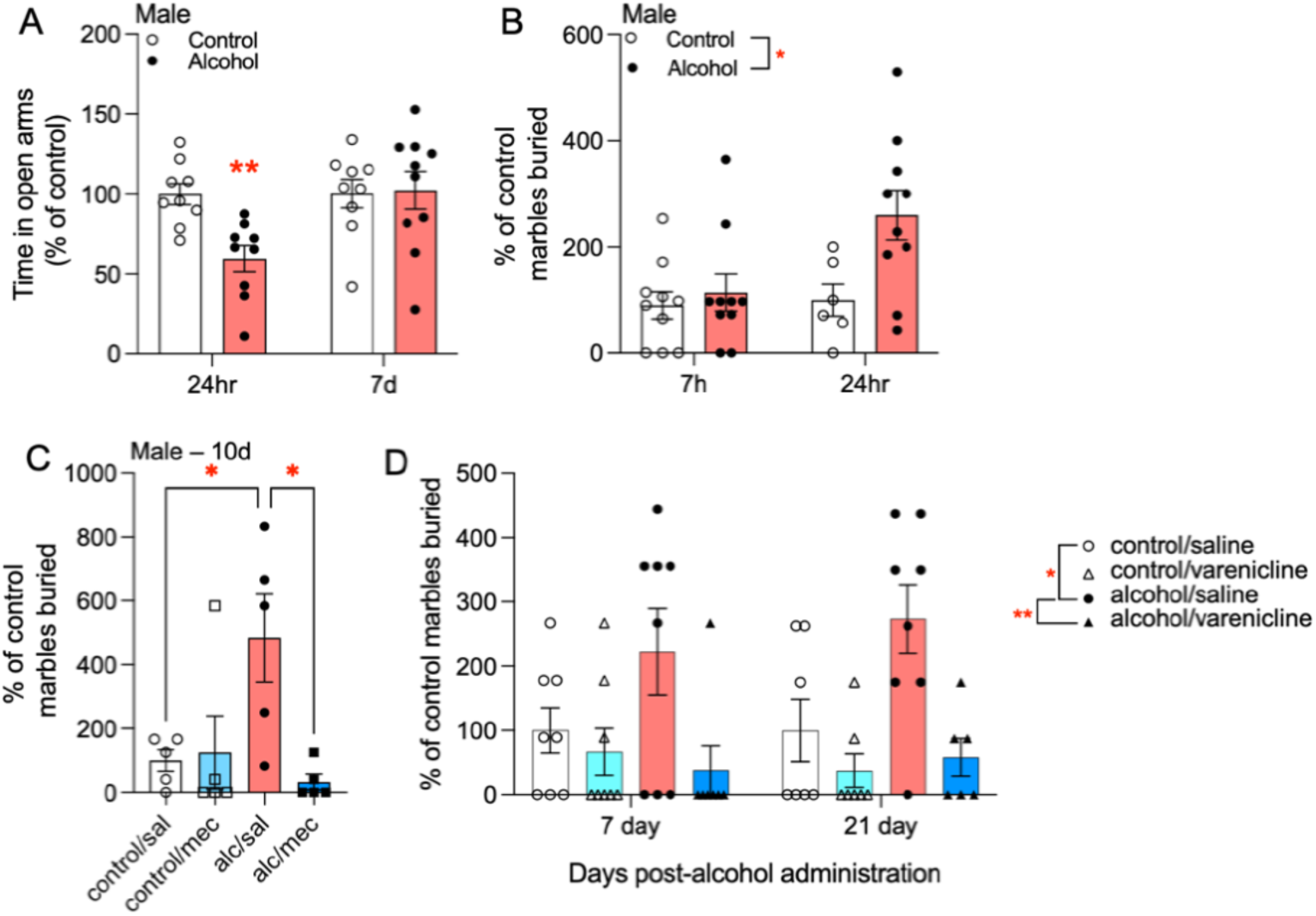
Affective behavior is perturbed during prolonged alcohol withdrawal and is attenuated by nicotinic acetylcholine receptor drugs in male mice. A) Alcohol withdrawn mice have decreased open arm time at 24h, but not at 7d, compared with control mice in the EZM. Sidak’s multiple comparisons test \*\**P*=0.009 compared with control mice, *n*=9-10 per group. B) Alcohol withdrawn mice show increased marble burying behavior over 24h compared with control mice. 2-way ANOVA main effect of group, \**P*=0.02, *n*=6-10 per group. C) The increase in marble burying behavior persists at 10d into alcohol withdrawal, which is attenuated by an *i.p.* injection of mecamylamine, a nAChR antagonist. Tukey’s multiple comparison test control/sal vs alcohol/sal, \**P*=0.04, alc/sal vs alc/mec \**P*=0.02, *n*=5 per group. D) Marble burying behavior disturbances persist to 21d into alcohol withdrawal and is attenuated by *i.p.* injections of varenicline. Repeated measures mixed effect model, main effect of group. Tukey’s multiple comparisons test control+4MP/saline vs alcohol+4MP/saline \**P*=0.015, alcohol+4MP/saline vs alcohol+4MP/varenicline \*\**P*=0.001, *n*=6-8 per group. Data shown as mean ± SEM.

We also assessed aspects of anxiety- and compulsive-like behavior with the marble-burying test (MBT) at 7h and 24h of alcohol withdrawal. There was a trend towards an interaction between treatment group and time point that was not significant, and significant main effects of treatment group and of time (2-way ANOVA, Finteraction (1, 32)=3.222, *P*=0.08; Ftime (1, 32)=4.242, *P*=0.048; Ftx group (1, 32)=5.946, *P*=0.02). The alcohol-treated mice showed greater marble-burying behavior, suggesting greater anxiety- and compulsive-like behavior, compared with the control-treated mice with the difference between treatment groups most prominent at the 24h withdrawal time point (**Fig. 2B**).

To determine the persistence of alcohol withdrawal-induced changes in the MBT, we tested additional time points into withdrawal. At 10d into withdrawal, alcohol-treated male mice still showed increased marble burying compared with control-treated mice (1-way ANOVA, F=4.823, *P*=0.01, Tukey’s multiple comparison test \**P*=0.04, alcohol+4MP/sal *vs.* control+4MP/sal, **Fig. 2C**). As nAChR drugs have shown efficacy in clinical trials of alcohol consumption [14–18] and we have previously shown that chronic alcohol administration increases the activity of cholinergic neurons of the brainstem of male mice [23], we tested whether mecamylamine, a nicotinic acetylcholine receptor (nAChR) antagonist, could attenuate the increased MBT behavior. We found that pre-treatment with 3 mg/kg *i.p.* mecamylamine completely attenuated the increase in marble-burying behavior in the alcohol-treated male mice compared with saline pre-treated mice (Tukey’s multiple comparison test \**P*=0.02, alcohol+4MP/sal *vs*. alcohol+4MP/mecamylamine, **Fig. 2C**).

We then tested varenicline, a partial agonist at α4β2 nAChRs, in the MBT at 7d and 21d into alcohol withdrawal in a separate cohort of mice. There was no interaction between time and treatment group nor a main effect of time, but there was a main effect of treatment group (mixed effects model with repeated measures for time, Finteraction (3,23)=0.3022, *P*=0.82; Ftime(1,23)=0.1118, *P*=0.74; Ftx group(3,28)=8.487, *P*=0.0004).

Tukey’s multiple comparisons test showed increased marble burying behavior at both 7d and 21d (\**P*=0.02 alcohol+4MP/saline *vs.* control+4MP/saline, **Fig. 2D**). Pre-treatment with 3 mg/kg *i.p.* varenicline significantly attenuated the increase in marble burying behavior at both time points compared with saline pre-treated mice (\*\**P*=0.001 alcohol+4MP/saline *vs.* alcohol+4MP/varenicline, **Fig. 2D**). Together, these data show that alcohol withdrawal-induced increases in anxiety- and compulsive-like marble burying behavior persist for at least 21d into withdrawal, and that mecamylamine and varenicline can attenuate this withdrawal-induced behavior in male mice.

### Female mice do not show somatic alcohol withdrawal signs or changes in EZM or MBT behavior

Female C57BL/6J mice received the same 9 daily injections of 2.5 g/kg alcohol plus 4MP or saline plus 4MP as the male mice. At 24h and 7d after the last injection, mice were assessed for somatic withdrawal signs, MBT and EZM. At 24h withdrawal, we did not observe an interaction between withdrawal sign and treatment group, nor a main effect of treatment group or withdrawal sign (Finteraction (3, 71)=0.3671, *P*=0.77; Fsign (3,71)=0.3671, *P*=0.77; Ftx group (1, 71)=0.091, *P*=0.76, **Fig. 3A**), indicating that female alcohol-treated mice did not show an increase in these somatic withdrawal signs compared with control-treated mice. We tested these female mice for anxiety-like behavior on the EZM at 24h withdrawal and found no differences in the time spent in the open arms between the alcohol- and control-treated mice (Student’s *t*-test *t*=0.0090, df=18, *P*=0.99, **Fig. 3B**). We also found no significant differences in the MBT between the alcohol- and control-treated mice at 24h withdrawal (Student’s *t*-test *t*=0.8645, df=16, *P*=0.40, **Fig. 3C**).

**Fig. 3.**
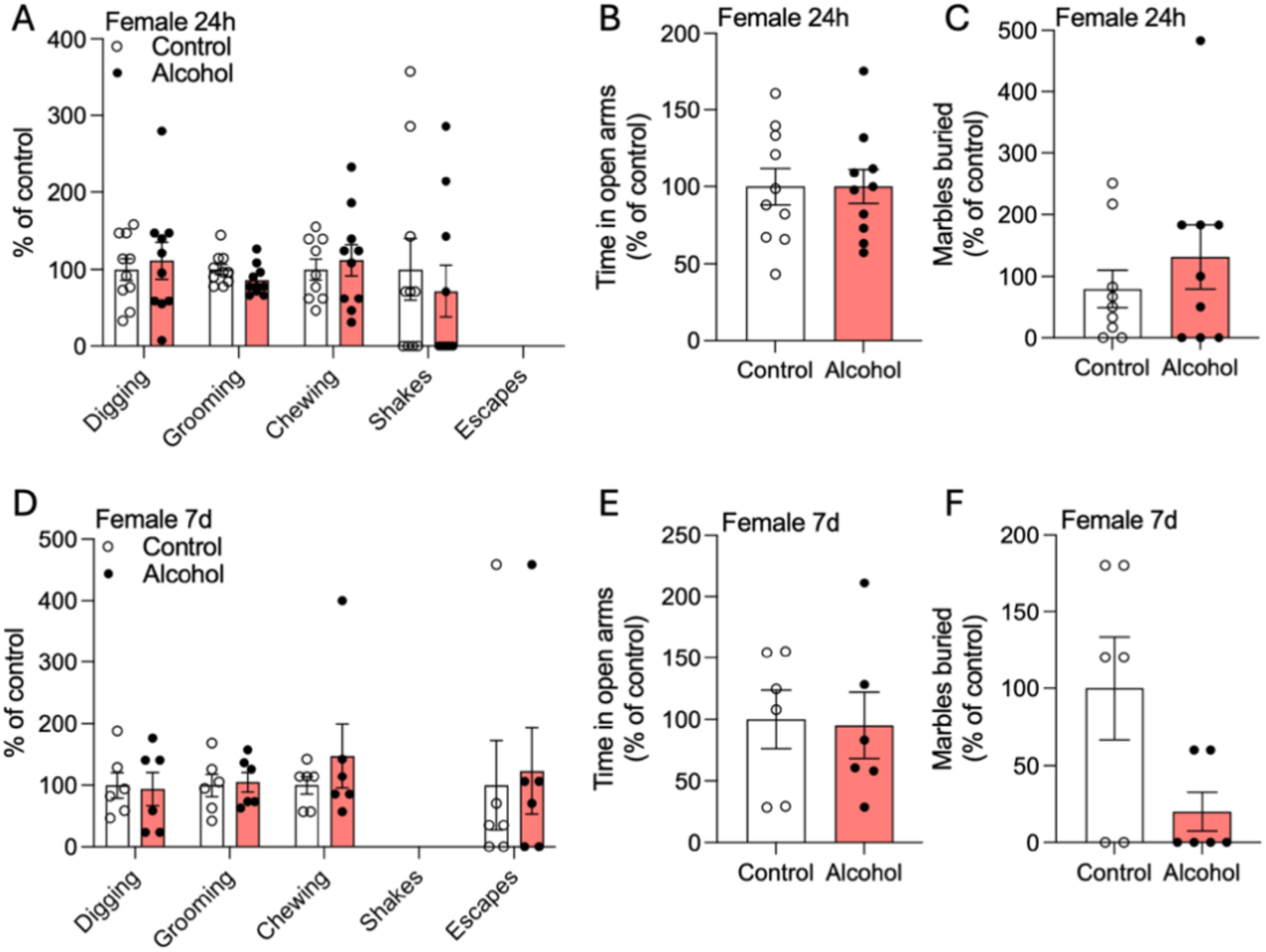
Female mice do not show alcohol withdrawal behaviors in the somatic withdrawal sign, elevated zero maze or marble burying assessments. Female alcohol withdrawal mice showed no statistically significant differences in A) somatic withdrawal signs, B) elevated zero maze or C) marble burying test at 24h of alcohol withdrawal compared with control mice. Zero escape attempts were recorded at 24h. There were no statistically significant differences in the D) somatic withdrawal signs, E) elevated zero maze or F) marble burying test at 7d withdrawal between the alcohol withdrawal mice compared with control mice. Zero shakes were recorded at 7d. The 24h and 7d were run in separate cohorts, *n*=9-10 per group for 24h, *n*=6 per group for 7d, data is shown as mean ± SEM.

We found similar results at the 7d withdrawal time point in a different cohort of female mice, with female alcohol-treated mice showing no significant increases in somatic withdrawal signs compared with control-treated mice (Finteraction (3, 40)=0.1473, *P*=0.93; Fsign (3, 40)=0.1474, *P*=0.93; Ftx group (1, 40)=0.3353, *P*=0.57, **Fig. 3D**).

Similarly, there was no significant difference between treatment groups in the EZM (Student’s *t*-test t=0.1348, df=10, *P*=0.90, **Fig. 3E**) or the MBT at the 7d withdrawal time point, but there was a trend for the female alcohol withdrawal mice to bury fewer marbles compared with control-treated female mice (Mann-Whitney test, *P*=0.11, **Fig. 3F**).

### Alcohol withdrawal produces a deficit in social interaction in female, but not male, mice

Alcohol withdrawal may produce different withdrawal signs in female mice compared with male mice, and negative affect may manifest differently in female mice compared with male mice. Therefore, we examined social behaviors with the social interaction test (SIT) after the 9-day alcohol administration procedure. Female mice were tested at 24h and 7d after the last injection in a social interaction chamber with a novel mouse of the same sex. There was no interaction between time point and treatment group nor a main effect of time, but there was a significant main effect of treatment group (mixed effect model with repeated measures for time, Finteraction (1,31)=1.699, *P*=0.20; Ftime (1, 31)=0.0011, *P*=0.92; Ftx group (1, 31)=4.442, \**P*=0.04).

Alcohol-treated female mice had a reduced interaction ratio compared with control-treated female mice, which was most prominent at the 24h withdrawal time point (**Fig. 4A**). The interaction ratio for the empty arena was not significantly different between the alcohol- and control-treated mice, indicating no effect of alcohol withdrawal on the exploration of the empty arena (2-way repeated measures ANOVA, Finteraction (1,17)=0.0001, *P*=0.99; Ftime (1,17)=0.4927, *P*=0.49; Ftx group (1,17)=1.127, *P*=0.30, **Fig. 4B**).

**Fig. 4.**
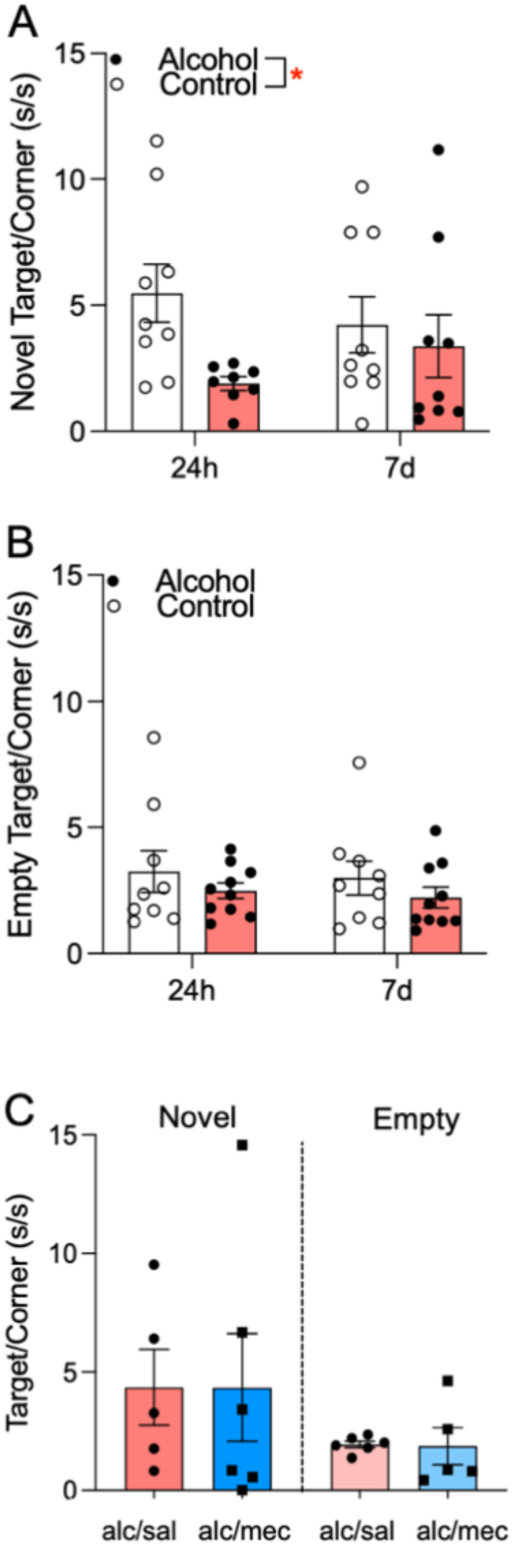
Female mice exhibit alcohol withdrawal-induced deficits in social interaction. A) Female mice show deficits in social interaction with a same-sex novel mouse at 24h and 7d of alcohol withdrawal. Repeated measures mixed effect model, main effect of group \**P*=0.04. B) Interaction with an empty social chamber was not different between alcohol withdrawn and control mice. *n*=8-9 per group. C) A separate group of alcohol withdrawn females were given *i.p.* saline or mecamylamine prior to the social interaction test at 24h withdrawal. There was no difference between the saline-and mecamylamine-treated groups for social interaction with the novel mouse (left side) or exploration of the empty social chamber (right side). *n*=5-6 per group, data shown as mean ± SEM.

We then tested whether mecamylamine could attenuate social interaction deficits in separate cohort of alcohol-treated female mice. Mice were pre-treated with 3 mg/kg *i.p.* mecamylamine 15 min prior to the social interaction test. There was no difference in the interaction score between mice that were pre-treated with mecamylamine versus saline in their interaction with a novel mouse (Welch’s corrected *t*-test, *t*=0.004057, df=8.553, *P*=0.99). There was also no effect of mecamylamine compared with saline on interaction with the empty arena (Welch’s corrected *t*-test, *t*=0.08608, df=4.248, *P*=0.94, **Fig. 4C**). These data indicate that antagonism of nAChRs with mecamylamine does not affect female social interaction behavior in alcohol withdrawal.

### Male mice do not show deficits in social interaction during alcohol withdrawal

We then returned to the male mice to determine if alcohol withdrawal produces deficits in SIT in males. Male mice were tested at 24h and 7d after the last injection. We found no interaction between time and treatment group, no main effect of treatment group, but a significant main effect of time (mixed effect model with repeated measures for time, Finteraction (1,13)=0.020, *P*=0.89; Ftime (1, 13)=15.72, \*\**P*=0.002; Ftx group (1, 14)=2.296, *P*=0.15). These results indicate that male mice in both treatment groups showed a reduced interaction score at the 7d time point compared with the 24h time point (**Fig. 5A**). There was no main effect of withdrawal time point or treatment group, nor an interaction between time point and treatment group for exploration of the empty social interaction chamber in male mice, indicating that alcohol withdrawal does not affect the exploration of the empty chamber (mixed effects model with repeated measures for time, Finteraction (1,27)=0.019, *P*=0.89; Ftime (1,27)=0.3316, *P*=0.57; Ftx group (1,27)=1.418, *P*=0.24, **Fig. 5B**). These data indicate that male mice do not show deficits in social interaction behavior in alcohol withdrawal.

**Fig. 5.**
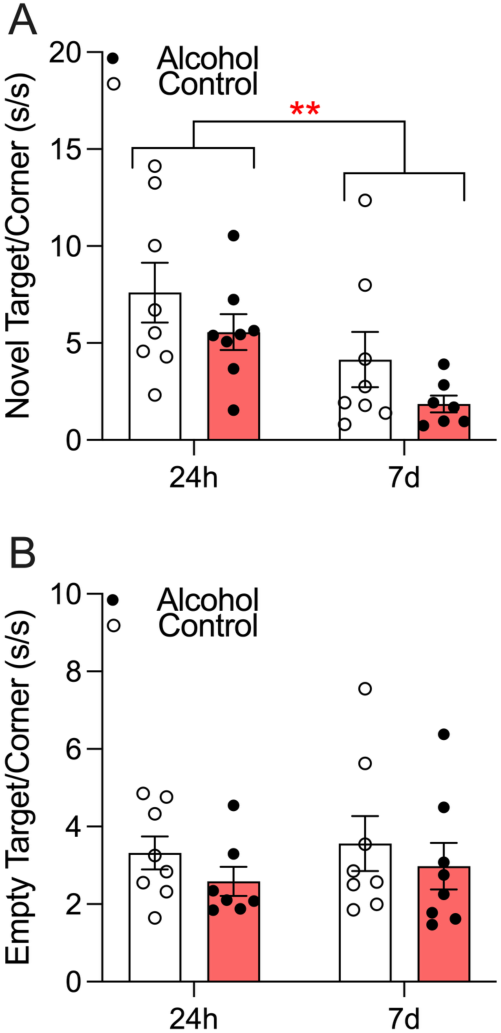
Male mice do not show deficits in social interaction during alcohol withdrawal. A) Male mice show decreased social interaction between 24h and 7d. Repeated measures mixed effect model, main effect of time \*\**P*=0.002. There was no difference between the control and alcohol-withdrawal groups for social interaction, main effect of group *P*=0.15. B) Interaction with an empty social chamber was not different between alcohol withdrawn and control mice. *n*=7-8 per group, data shown as mean ± SEM.

### Alcohol clearance does not differ between male and female mice

Although we administered 2.5 g/kg alcohol with 4MP to males and females, and thus controlled the amount of alcohol the mice received, we investigated the clearance of alcohol to determine whether sex differences in alcohol metabolism could contribute to the differences in alcohol withdrawal behaviors. Male and female mice were given one injection of 2.5 g/kg alcohol + 9 mg/kg 4MP (acute alcohol treatment group), or 9 daily injections of 2.5 g/kg alcohol + 9 mg/kg 4MP (chronic alcohol treatment group).

Blood samples were taken at 15, 45 and 75 minutes after the last injection. We found no significant 3-way interaction between length of alcohol treatment (acute vs. chronic), time point or sex (3-way ANOVA, Falcohol gpXtimeXsex (2,61)=0.2342, *P*=0.79) and no significant 2-way interactions (Falcohol gpXsex (1, 61)=0.2629, *P*=0.61; FtimeXsex (2, 61)=0.5446, *P*=0.58; Falcohol gpXtime (2, 61)=2.473, *P*=0.09). There were also no significant main effects of time or sex (F time(2, 61)=1.552, *P*=0.22; Fsex (1,61)=2.014, P=0.16); however, there was a strong trend for a main effect of the length of alcohol treatment (Falcohol gp (1, 61)=3.828, *P*=0.055, **Fig. 6A**). As sex was not significant, we collapsed males and females and analyzed the data as a 2-way ANOVA to gain further insight into alcohol clearance over time between the acute versus chronic alcohol-treated groups.

**Fig. 6.**
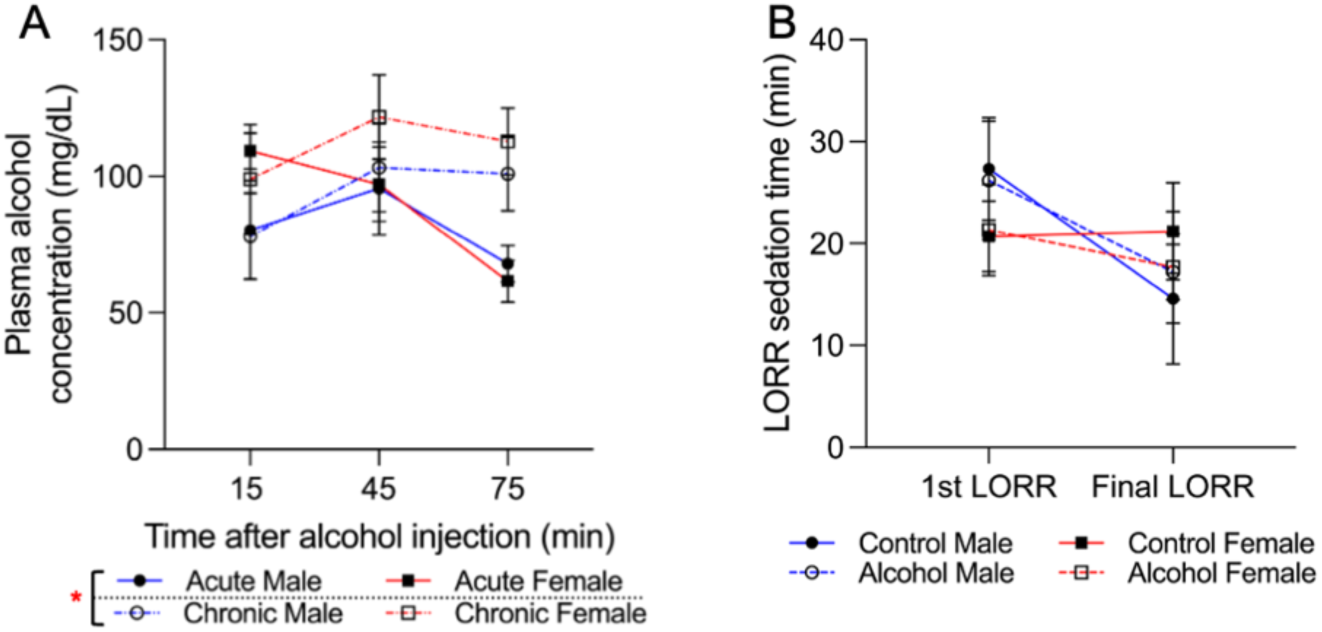
Alcohol clearance and sedation are not different between male and female C57BL/6J mice. A) Alcohol clearance after a single injection of 2.5 g/kg alcohol + 9 mg/kg 4MP (acute), or after 9 daily injections of 2.5 g/kg alcohol + 9 mg/kg 4MP (chronic), was not different between males and females. Three-way mixed effects model did not show a main effect of sex, *P*=0.16. Collapsing sex resulted in a main effect of treatment between acute and chronic alcohol administration, \**P*=0.04. Acute alcohol group *n*=5 per sex, chronic alcohol group *n*=7-8 per sex. B) Alcohol sedation time after a 4.0 g/kg *i.p.* injection was measured before (1^st^ LORR) and after (final LORR) 9d of 2.5 g/kg + 9 mg/kg 4MP (alcohol group) or saline + 9 mg/kg 4MP (control group) in males and female mice. Mixed effects model showed no significant 3-way or 2-way interactions between sex, time and treatment group, and no main effects of sex, time or treatment. *n*=5-7 per sex per group. Data shown as mean ± SEM.

We found no significant interaction between the alcohol treatment group and time, nor a main effect of time, but there was a significant main effect of alcohol treatment group (acute *vs.* chronic) (Falcohol gpXtime (2, 67)=2.650, *P*=0.78; Ftime (2, 67)=1.603, *P*=0.21; Falcohol gp (1, 67)=4.186, \**P*=0.045). These data suggest that alcohol clearance is slower in male and female mice that have been chronically treated with 9 days of 2.5 g/kg plus 4MP compared with mice that received only one injection. As there was no sex difference in alcohol clearance, this indicates that the differences we observed in somatic withdrawal signs, affective behavior and social interaction cannot be attributed to sex differences in alcohol clearance.

### Alcohol sedation does not differ between male and female mice

We assessed whether the sedative effect of a large dose of alcohol differed between male and female mice before and after the 9 days of alcohol+4MP treatment. Mice were assessed for alcohol sedation using the loss-of-righting-reflex (LORR) test with 4 g/kg *i.p.* alcohol. Mice were given a first LORR assessment, followed by 9 daily injections of 2.5 g/kg+4MP or saline+4MP, and assessed for LORR again 24h after the last injection. There was no significant 3-way interaction between LORR session (1^st^ *vs.* final LORR), sex or treatment group (FLORR sessionXtx groupXsex (1,7)=0.2943, *P*=0.60), nor any significant 2-way interactions (Ftx groupXsex (1, 11)=0.02703, *P*=0.87; FLORR sessionXtx group (1, 11)=0.02473, *P*=0.88; FLORR sessionXsex (1, 7)=2.896, *P*=0.13). There was no main effect for LORR session, sex or treatment group (FLORR session (1, 11)=2.168, *P*=0.17; Fsex (1, 11)=0.05479, *P*=0.82; Ftx group (1, 11)=0.06217, *P*=0.81; **Fig. 6B**). These data indicate that the sedative effect of 4 g/kg alcohol does not significantly differ due to sex, alcohol+4MP treatment or time.

## Discussion

Early alcohol withdrawal in humans consists of somatic withdrawal signs along with increases in negative affect. The somatic withdrawal signs abate over time; however, the increased negative affect, which includes irritability, anxiety and anhedonia, can persist for months or even years [6,7,24]. This prolonged withdrawal phase is critically important as the persistent negative affect has long been recognized as a risk factor for relapse back to alcohol use [6,8]. Moreover, there is a lack of targeted pharmacotherapy that addresses the increased negative affect during prolonged withdrawal [5,7,24].

Alcohol withdrawal in rodents has been achieved using multiple procedures including forced alcohol vapor administration, systemic alcohol injections, and voluntary continuous, intermittent and binge alcohol drinking procedures [11,25]. Voluntary alcohol consumption most closely models the human experience of alcohol drinking and allows for inter-individual variation in drinking levels. However, in C57BL/6J mice, the most commonly used mouse strain in alcohol studies, female mice consistently demonstrate greater alcohol consumption on a g/kg basis compared with male mice, which we have also observed in our studies [26–28]. In this study, we ensured that all mice received the same dose of alcohol per body weight and used 9 daily *i.p.* injections of 2.5 g/kg alcohol paired with 4MP, an alcohol dehydrogenase inhibitor [29]. This procedure was adapted from a prior study by Perez and colleagues that used a similar injection protocol and showed that it produced spontaneous withdrawal signs [29], increased anxiety-like behavior in the open field test and elevated zero maze, and increased marble burying behavior at 24h withdrawal that was equivalent to the withdrawal signs observed after a liquid diet of alcohol for 6 weeks [25].

We found that somatic alcohol withdrawal signs changed over time, reflecting a progression in the manifestation of alcohol withdrawal in male C57BL/6J mice. There was increased shaking behavior at 7h of withdrawal that coincides with the time of peak handling-induced convulsions in male mice [28], which is often used as a measure of alcohol withdrawal-induced hyperexcitability and seizure susceptibility. At 24h, the increased shaking had returned to baseline and there was an increase in chewing behavior. At 7d, we observed only increased digging behavior, and this disturbance in natural digging behavior may also be reflected in the increased marble burying behavior during alcohol withdrawal in male mice. Alcohol withdrawal-induced negative affect also showed changes over time, with an increase in anxiety-like behavior at 24h that returned to baseline at 7d, and increased anxiety- and compulsive-like behavior in the MBT that peaked at 24h and remained elevated at 7, 10 and 21d into withdrawal. These temporal changes in affective alcohol withdrawal signs demonstrate that different aspects of negative affect emerge over the course of alcohol withdrawal.

In contrast to male mice, we found that female C57BL/6J mice administered the same dose of alcohol using the same procedure did not show somatic withdrawal signs at 24h of withdrawal. We found that female mice also did not show significant increases in anxiety- and/or compulsive-like behavior when assessed on the EZM or the MBT, whereas male mice showed behavioral disturbances in both assays at 24h. A prior study examining both sexes showed that 36h withdrawal from voluntary alcohol consumption produced increased marble burying behavior in male but not in female mice, even though female mice consumed significantly more alcohol than the male mice [30]. Other studies have shown no changes in anxiety-like behavior in female mice in the elevated plus maze at 14d withdrawal [31,32]. Alcohol withdrawal-induced anhedonia assessed using the forced swim test and sucrose preference test showed that male, but not female, mice exhibited increased depressive-like behavior at 3 weeks withdrawal, but both sexes showed a deficit in sucrose preference [33].

Historically, female rodents have been included in preclinical studies at far lower rates [34]. Alcohol withdrawal studies that do include female mice have shown reduced somatic withdrawal signs and no changes in negative affective behavior in females (for review see [11]). Part of the discrepancy may be due to the behavioral tests used to assess negative affect, which have largely been developed using male mice and exploit the conflict between the drive to explore a novel space compared with a safer environment. Female animals may exhibit different negative affective behaviors and alcohol withdrawal may manifest differently in female mice compared with male mice. To assess behavior that may be more relevant to negative affect in female mice, we examined the effect of alcohol withdrawal on social interaction and found a withdrawal-induced deficit that was most prominent at 24h. We then examined male mice and found no significant effect of withdrawal on social interaction in male mice, indicating a sex difference in withdrawal-induced effects on this behavior.

There was no significant difference in alcohol clearance between sexes, indicating that the differences in withdrawal signs cannot be attributed to differences in alcohol clearance. Interestingly, we found that chronic alcohol plus 4MP administration resulted in reduced alcohol clearance compared with a single administration of alcohol plus 4MP in both sexes. Chronic alcohol administration in humans and rodents increases alcohol metabolism through the induction of cytochrome P450 2E1 expression [35], therefore we postulate that the reduction in alcohol clearance after chronic alcohol+4MP in our study is due to an accumulating effect of alcohol dehydrogenase inhibition.

Overall, we show that alcohol withdrawal manifests differently in male versus female mice, and that withdrawal-induced negative affect in female C57BL/6J mice may be better captured by tests of social interaction. Female mice may also exhibit more subtle somatic withdrawal signs compared with male mice and future work to delineate these signs would be valuable to better understand how alcohol withdrawal affects female mice.

The persistent negative affect during prolonged withdrawal in male mice likely reflects underlying persistent adaptations in neurobiology caused by alcohol and/or withdrawal from alcohol. We showed that pre-treatment with varenicline and mecamylamine were both able to attenuate the increased marble burying behavior in prolonged alcohol withdrawal in male mice. Mecamylamine is a non-specific nicotinic acetylcholine receptor (nAChR) antagonist that affects all nAChR subtypes, whereas varenicline is a partial agonist at the α4β2 and α3-containing nAChR subtypes [36,37], and a full agonist at α7 nAChRs [38]. That both mecamylamine and varenicline are able to attenuate the withdrawal-induced marble burying behavior suggests that this long-lasting increase in negative affect is mediated by sustained and elevated cholinergic activity at nAChRs in male mice.

Prior pre-clinical research on cholinergic signaling and nAChRs have focused on the role of nAChRs in mediating alcohol reward. Alcohol consumption increases extracellular acetylcholine (ACh) levels in the ventral tegmental area (VTA) in male Wistar rats [39]. Mecamylamine [40] and varenicline can reduce alcohol consumption in a binge paradigm in male C57BL/6J mice, and the α4 nAChR subunit in the VTA is necessary for varenicline to reduce consumption in male mice [41,42]. The contribution of cholinergic signaling and nAChR expression, particularly outside of the mesolimbic circuitry, to alcohol withdrawal is far less studied. We recently showed that acute and chronic alcohol administration increases cholinergic neuron activation, via increased c-Fos protein and transcript expression, in the pedunculopontine tegmentum (PPN) of male mice [23]. The PPN and its neighboring cholinergic nucleus, the laterodorsal tegmentum (LDT), are brainstem regions that provide broad cholinergic innervation to the midbrain, hindbrain and the spinal cord [43]. We showed the chronic injections of 2.0 g/kg alcohol for 15 days increased cholinergic neuron activation in the PPN, but not the LDT, when measured 2h after the last alcohol injection [23]. Our current focus is to determine whether this increase in cholinergic neuron activity in the PPN is sustained into alcohol withdrawal, and we postulate that mecamylamine and varenicline act to reduce negative affect by attenuating the increased cholinergic signaling in the brainstem.

In contrast to the males, our prior work showed that acute and chronic alcohol administration did not increase cholinergic neuron activity in the PPN or LDT in female mice [23]. Here, we found that mecamylamine did not affect social interaction behavior in the female alcohol-treated mice. Together, our data suggest that alcohol administration and alcohol withdrawal produces a sustained increase in cholinergic activity that mediates withdrawal-induced negative affect in male, but not in female, mice. Thus, the neurobiological changes caused by alcohol withdrawal and the mechanisms that mediate withdrawal-induced negative affect differ by sex, which has important implications in the development of AUD pharmacotherapies for alcohol withdrawal. Varenicline is a FDA-approved for smoking cessation [13] that is also in clinical trials for AUD. Varenicline has shown promise in reducing alcohol consumption and craving in multiple clinical trials [14–18]. However, nearly all clinical trials investigating varenicline have more male than female participants. Critically, a clinical trial that examined sex differences found that varenicline was more effective at decreasing heavy drinking in men compared with women [44]. Varenicline has not been investigated for the reduction of negative affect in prolonged alcohol withdrawal. These results emphasize the importance of studying both males and females, and suggest that varenicline may be a promising therapeutic for treatment of prolonged alcohol withdrawal in men, but may not be as effective in women.

## Data availability statement

The datasets generated during and/or analyzed during the current study are available from the corresponding author on reasonable request.

## Acknowledgements

This work was supported by R25 DA057802, P30 DA048742 and a University of Minnesota Foundation Award.

## Author contributions

N.R.P., S.M.S., M.S., R.F.D., S.E. and A.M.L. collected the data, N.R.F., S.M.S., S.E. and A.M.L. analyzed the data, and N.R.P., S.M.S. and A.M.L. wrote and edited the manuscript.

## Competing interests

The authors declare no competing interests.

## Notes

### Competing Interest Statement

The authors have declared no competing interest.

